# Malaria Outbreak Detection with Machine Learning Methods

**DOI:** 10.1101/2020.07.21.214213

**Authors:** Gurcan Comert, Negash Begashaw, Ayse Turhan-Comert

**Affiliations:** Department of Comp. Sc., Physics, and Engineering, Benedict College, 1600 Harden St., Columbia, SC 29204

**Keywords:** Malaria outbreak detection, random decision trees, logistic regression, Gaussian processes

## Abstract

In this paper, we utilized and compared selected machine learning techniques to detect malaria outbreak using observed variables of maximum temperature, minimum temperature, humidity, rainfall amount, positive case, and Plasmodium Falciparum rate. Random decision tree, logistic regression, and Gaussian processes are specially analyzed and adopted to be applied for malaria outbreak detection. The problem is a binary classification with outcomes of outbreak or no outbreak. Sample data provided in the literature from Maharashtra, India is used. Performance of the models are compared with the results from similar studies. Based on the sample data used, we were able to detect the malaria outbreak without any false positive or false negative errors in the testing dataset.

## 1 Introduction

Malaria outbreak detection is an important problem impacting millions of lives annually. Early prediction of outbreak helps health organizations to take actions and control the outbreak ([1]). There are several applications of machine learning methods in the literature. We focused on relevant studies and investigated them in details.

In the work of Vijeta Sharma et al., [1], support vector machine outperformed artificial neural networks. In this fundamental work, Support Vector Machine (SVM) and Artificial Neural Network (ANN) were used for Malaria prediction using a dataset from Maharashtra state. Authors showed that SVM was able to miss less outbreak detections than NN. Root mean squared errors are reported as 0.12 and 0.47 and correct detection rates were 89% and 77% for SVM and NN, respectively. Noor et al. [2] used medical intelligence, reported case incidence, and extreme climatic conditions to define risky locations. Authors geocoded Plasmodium Falciparum (pF) parasite rate across 49 endemic countries and territories in Africa from surveys undertaken since 1980. The data were used within a Bayesian space–time geostatistical framework to predict for 2000 and 2010 at a 1 squared kilometer (*km*^2^) spatial resolution. Population distribution maps were also integrated for risk quantification. They have assembled the largest geocoded repository of malaria infection prevalence data for Africa as part of an 8-year data search. They used these data to predict in space (1 *km*^2^ resolution) and time (2000 and 2010) the intensity of pF transmission across malaria-endemic countries and territories. Leopord Kakizimana et al. [3] used very similar batch of methods as in our study. The authors aimed to test hybrid classification and regression models to predict the disease outbreak using datasets where it has been observed that some of the single data mining techniques have accuracy weakness. They showed that the hybridization model should overcome the single model’s weakness by combining more than one technique where decision tree, random forest, Naïve Bayes multinomial, simple logistic, and Bayesian logistic regression were combined to achieve a hybrid based classification and regression model for predicting the disease outbreak with high accuracy. The authors also used the dataset from [1]. Their accuracy levels are close to ours in training, however, their validation accuracy is about 75% which is much lower than our validation accuracy. Babagana Modu et al. [4] used the partial least squares path modelling methodology to analyse the causal relationships among meteorological variables, e.g., minimum average temperature, maximum average temperature, relative humidity, wind speed, precipitation and solar radiation, and explored their impact on the outbreak of malaria. They used a total of 85,627 confirmed diagnosed cases of malaria incidence for a period of five years from 2009 to 2013. The distributional pattern of malaria cases reported in the study area shows an indication of high malaria incidence. Authors applied several machine learning algorithms, including Support Vector Machine (SVM), K-Nearest Neighbours (KNN), Naive Bayes and Decision Trees, to find the best predicting algorithm from the scikit framework [45] in Python. They reported 99 % detection accuracy with SVM, 75 % with logistic regression, 64 % with decision tree, and 81 % with KNN. Julius Ssempiira et al. [5] applied Bayesian spatio-temporal negative binomial models that were fitted on district-aggregated monthly malaria cases, reported by age groups. Weather data included rainfall, day and night land surface temperatures, normalized vegetation index, altitude, land cover, and distance to water bodies. The authors estimated the effects of climatic changes on the malaria incidence between 2013 and 2017 by modeling time varying climatic conditions, adjusting for the effects of intervention coverage, and socio-economic factors. Random effects at district level were used to model spatial correlation. Temporal correlation was taken into account by monthly random effects modeled by autoregressive processes. Models were also adjusted for seasonality. The study provides results on posterior estimates, but, it did not provide any accuracy or comparison to other methods.

From existing studies, varying accuracies were reported. As interesting classification problem, we analyzed the impact of different variables and adopted random decision trees, logistic regression, and Gaussian processes to predict malaria outbreak. We used the data presented in ([1]) and compared the results in our summary table. Based on the test data used, our results show 100% accuracy in prediction. The rest of the paper is organized as follows. In 2, we define the problem and discuss the data used in this study. In section 3, we discuss the methods and training in detail. In section 4 we present our results. Finally, in section 5, we summarize and discuss our findings.

## 2 Problem Definition

Given a set of observed variables **X** = (*X*_1_, *X*_2_*, …, X_n_*), we first aim to understand which of these variables correlate highly with the occurrence of the response variable. This is in fact a problem of dimensional reduction or also known as principal component analysis. In a basic classification algorithm, we observe a variable that we assume to be able infer about its latent states, i.e., probability of response variable *Y* is being at latent state *Z* = *z* given observed *X* variable, *P* (*Z* = *z*|*X*). However, not all problems can easily or without error can be set up this way. If we think of the problem as a simple regression problem, the observed variables may not be able to map the variation in Y perfectly.

For instance, in Fig. 1, we can see that it is not easy to determine what would be a combination of variables used to map to a binary outbreak variable. Thus, our selection of methods can be explained intuitively as we are looking for methods which can incorporate such unobserved variations into account. We use correlations in the case of Gaussian Process, regression set up and completing as choices in the case of logistic regression, and bootstrapping in the case of random decision trees.

**Figure 1:**
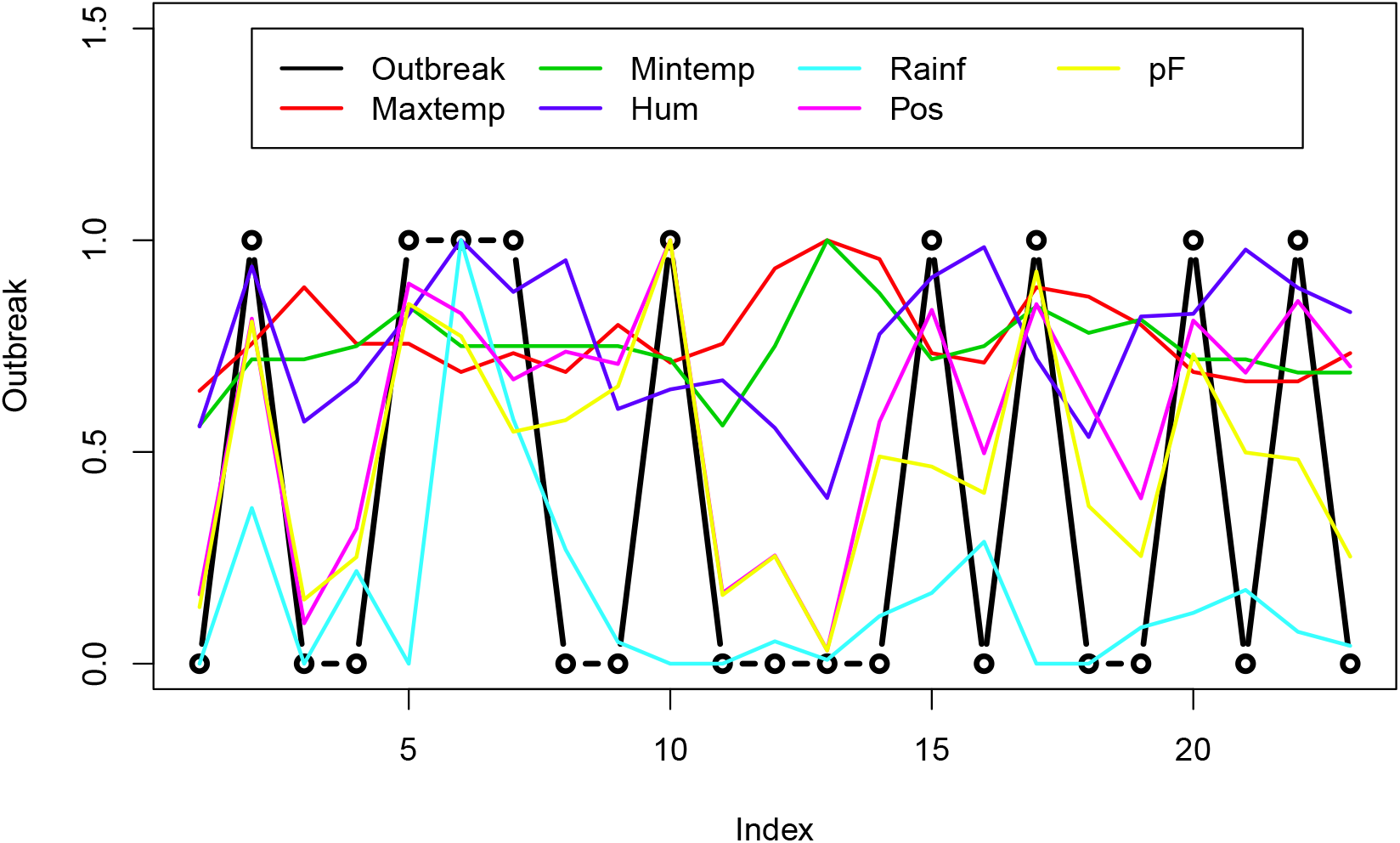
Behavior of training data with respect to outbreak

### 2.1 Data Description

We are using the Malaria data samples provided in [1]. Sample of the data contains 38 rows of observations. The variables considered are average monthly maximum temperature (in degree Celcius), minimum temperature (in degree Celcius), relative humidity (%), rainfall amount (in inches), total number of positive cases, total number of Plasmodium Falciparum(pF) cases, and response of outbreak (1) or no outbreak (0). We used first 23 rows of data (see Fig. 1) to train our models and utilized the remaining 15 rows of data for testing the performance of the proposed models.

Figure 2 shows correlation matrix of the variables. Simply we see higher to lower correlations between outbreak and pF, positive case, humidity, maximum temperature, and rain fall, respectively. Logically, we can start including positive cases and pF in our models then include others humidity, maximum temperature, and rainfall in order.

**Figure 2:**
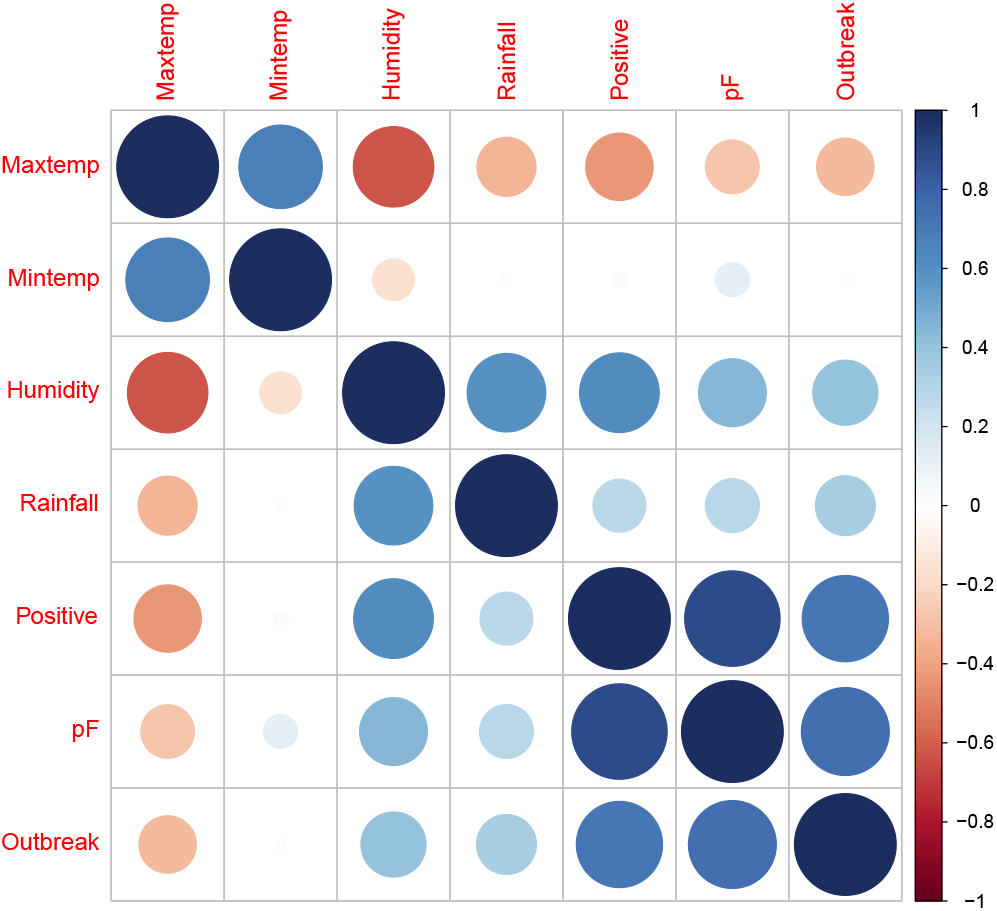
Correlation matrix of training data with respect to outbreak

## 3 Methodology

In this section, we present the technical details of the three proposed machine learning methods used for detecting malaria outbreak.

### 3.1 Decision Trees

Decision trees (DTs) are nonparametric decision rules to regress multivariate input of observations to response variables. Using main algorithms (e.g., ID3 (Iterative Dichotomiser 3) and CART (Classification And Regression Tree)), trees can be learned recursively. After determining the root either with correlation or Gini factor, recursively thresholds are found from the training data until reaching to a leaf which is practically a prediction of a response value. In this study, we formulated the following recursive algorithm (algorithm 1) in *R* programming.

#### Algorithm 1 DT learning algorithm

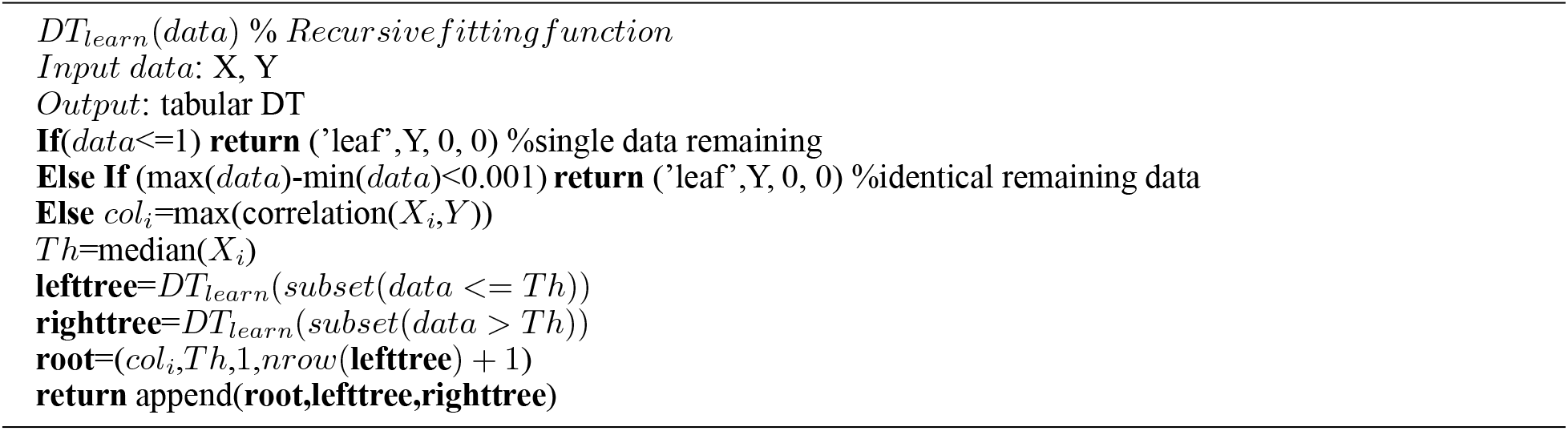

After training the tree, we can use it for prediction using the function in 2. This is essentially reading the DT table generated by *DT*_*learn*_ function in 1.

#### Algorithm 2 DT learning algorithm

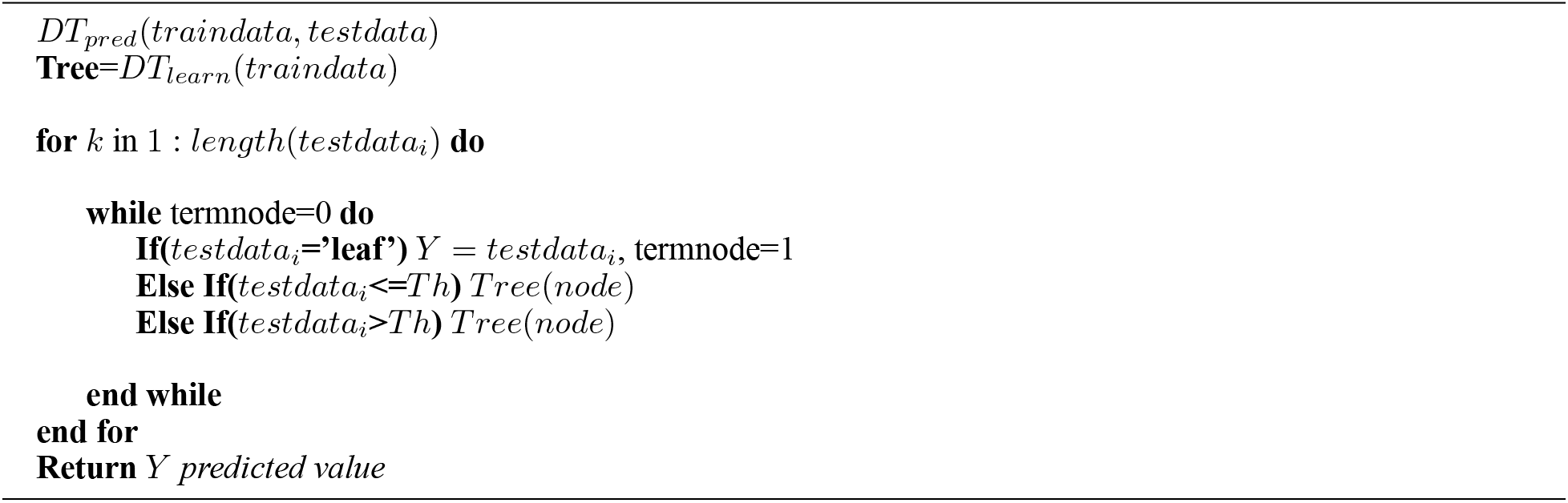

Note that, we can either train the decision tree once or we can randomly select from the training samples and derive multiple decision trees (i.e., random trees). In this study, we used random trees and took average at the end to make decision. Figure 3 shows one of the trained random decision trees. The Root is the first node and in the example figure *pF* ≤ 387.8 results as no outbreak. Otherwise, we look other threshold values to be able to reach a leaf. Denoted by green nodes, the leaves are the nodes for Y predictions (*Y* = 1 outbreak, *Y* = 0 no outbreak). After 1,000 decision trees, we took the average for the predictions 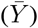. In a way, we equally weighted 1000 DTs and made decision 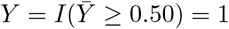, otherwise *Y* = 0.

**Figure 3:**
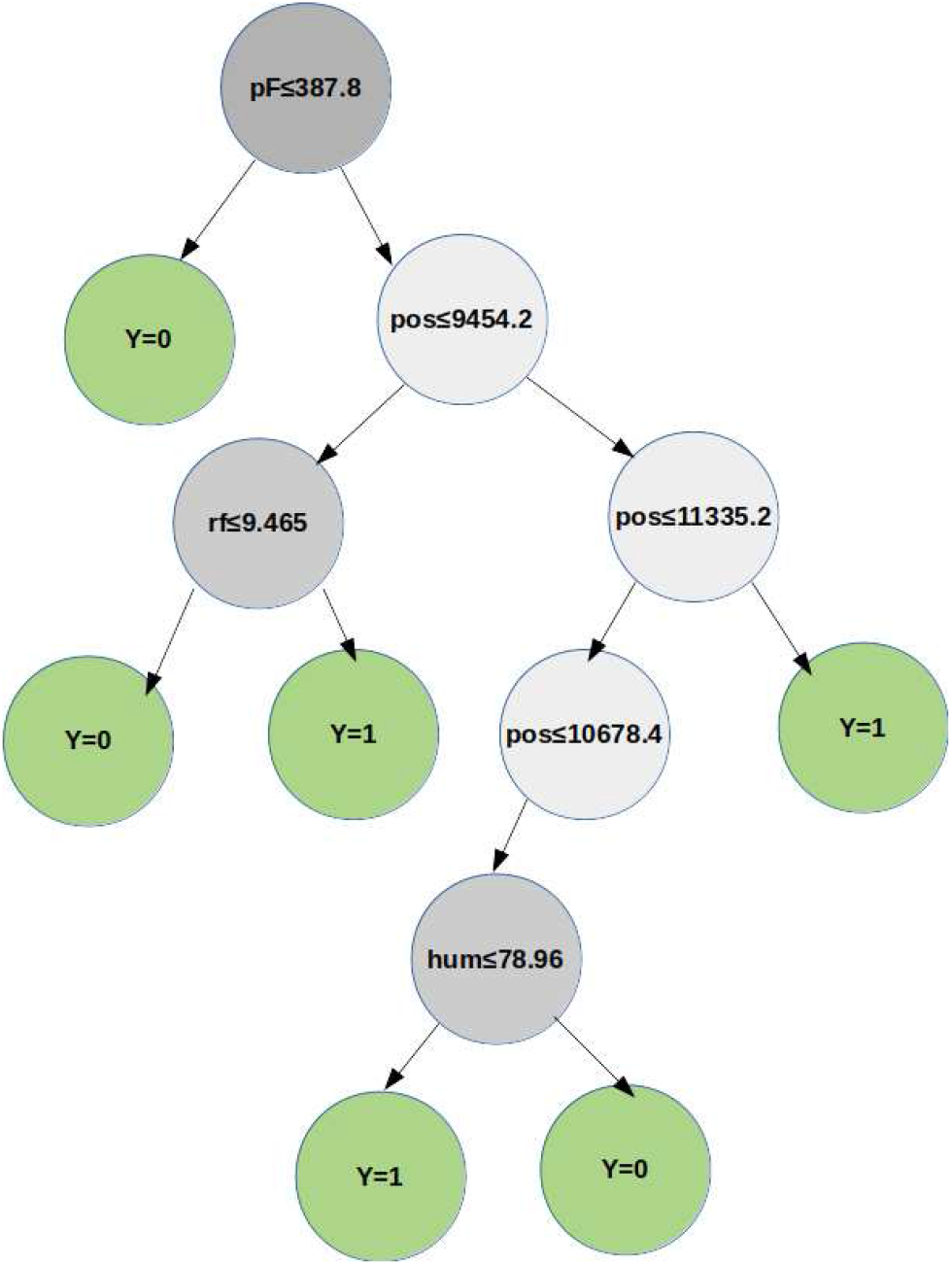
A sample of the trained random decision trees

**Figure 4:**
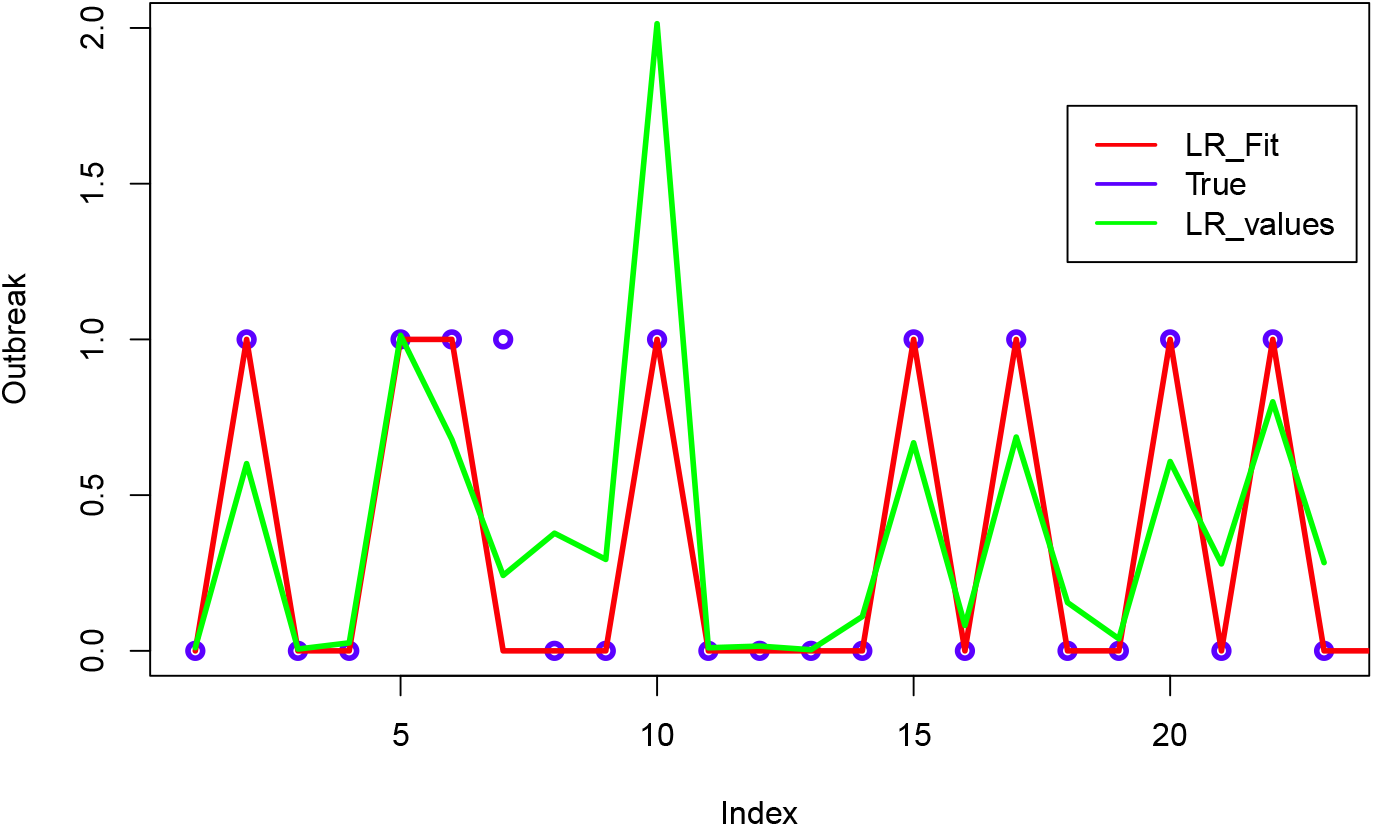
Fitting results from logistic regression

### 3.2 Logistic Regression

Next, we fitted multiple logistic regression models with different distribution families given in base *R* functions under generalized linear models *glm*. The best performing model is found to be either Poisson or quasi-Poisson family. The difference is that quasi-Poisson uses quasi-likelihood for parameter estimation and is more appropriate with overdispersion. The models provide probabilities as response. We determine the outcome based on 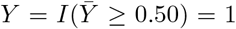, otherwise *Y* = 0. After checking other combinations, only maxtemp, rf, pos, and pF are used in the model fit. These are consistent with the DT variables. Like in DT, we did not transform or normalize the variables. However, we had to determine important variables and reduce the variables based on performances. The model parameters are intercept *β*_0_ = −5.142, maxtemp (*X*_1_) *β*_1_ = −0.714, pos (*X*_2_) *β*_2_ = 6.234, and pF (*X*_3_) *β*_3_ = 0.116.

Suppose that *Y* = *y*_1_*, y*_2_*, …, y_n_* are Poisson with *P* (*λ*). Log-linear version of a Poisson regression (logistic regression with Poisson family) can be written as in Eq. (1).

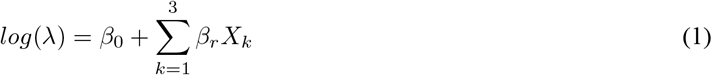

where, the model parameters are *β*_0_, …, *β*_4_ and the variables are *X*_*k*_s.

### 3.3 Gaussian Processes

Finally, we used Gaussian Processes as natural alternatives with correlation kernel possibly dealing with spatial and temporal correlations as well as collinearities. Gaussian process can be described as regression on function space where over infinitely many dimensions a data generation process is explained via mean *μ*(*x*) and covariance functions *K*(*x*) in a semi-parametric way. The only parameters are those in the covariance functions which describe the magnitude and length of the volatility. We applied GP using *tgp*, *GPfit*, and *mlegp* packages in *R* from [6, 7, 8], respectively. After carefully adopting and selecting parameters for our problem, we obtained similar results. For GPs, we made a transformation of the variables by dividing each variable *X*_*i*_ by *M* = max{*X*_*i*_ : *i* = 1, · · · *n*}. The transformed variables will all be in [0, 1]^*d*^. Only maxtemp, rf, and pos variables were used in model fit.

We adopted the GP notations from [7]. Let **X** = (*x*_*i*_1, *x*_*i*_2, …, *x*_*i*_*d*)^*T*^ denote input observations (*i* = 1, *…*, 23, *n* = 23 in the numerical examples) with dimension *d* = 3. Output vector or outbreak results are *Y* =*y*(**X**)=(*y*1, …, *y*_*n*_)^*T*^ for *y*_*i*_=*y*(*x*_*i*_). We can express *y*(*x*_*i*_)=*μ*+*z*(*x*_*i*_) with *μ* is overall intercept and we model *z*(*x*_*i*_) with 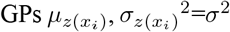 and covariance function *Cov*(*z*(*x*_*i*_), *z*(*x*_*j*_))=*σ*^2^*R*_*ij*_ . Out of several kernels or correlation structures (exponential, linear, and Matérn), we found that

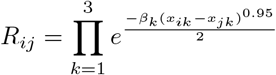

for all *i, j*.

GP fitting minimizes negative log marginal likelihood with respect to hyperparameters and noise level. From our model fit, hyper parameters were calculated. Power of the exponential correlation function was selected 0.95. Correlation parameters were estimated as *β*_1_ = −9.802, *β*_1_ = −0.026, and *β*_3_ = 0.067, and *σ*^2^ = 0.273.

## 4 Numerical Results

In this section, we present the results from the methods utilized. Figure 6 show the true values between samples 24-38 in [1, 3], average values are shown as *DT*_*values*_, and detection results are after treating the values with 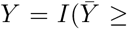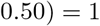. As it can be seen from the figure, we are able to correctly detect the outbreak using all 6 variables with any transformation or reducing the dimensionality of the variable set. Decision trees are able to handle this automatically while determining the significant factor and the threshold.

**Figure 5:**
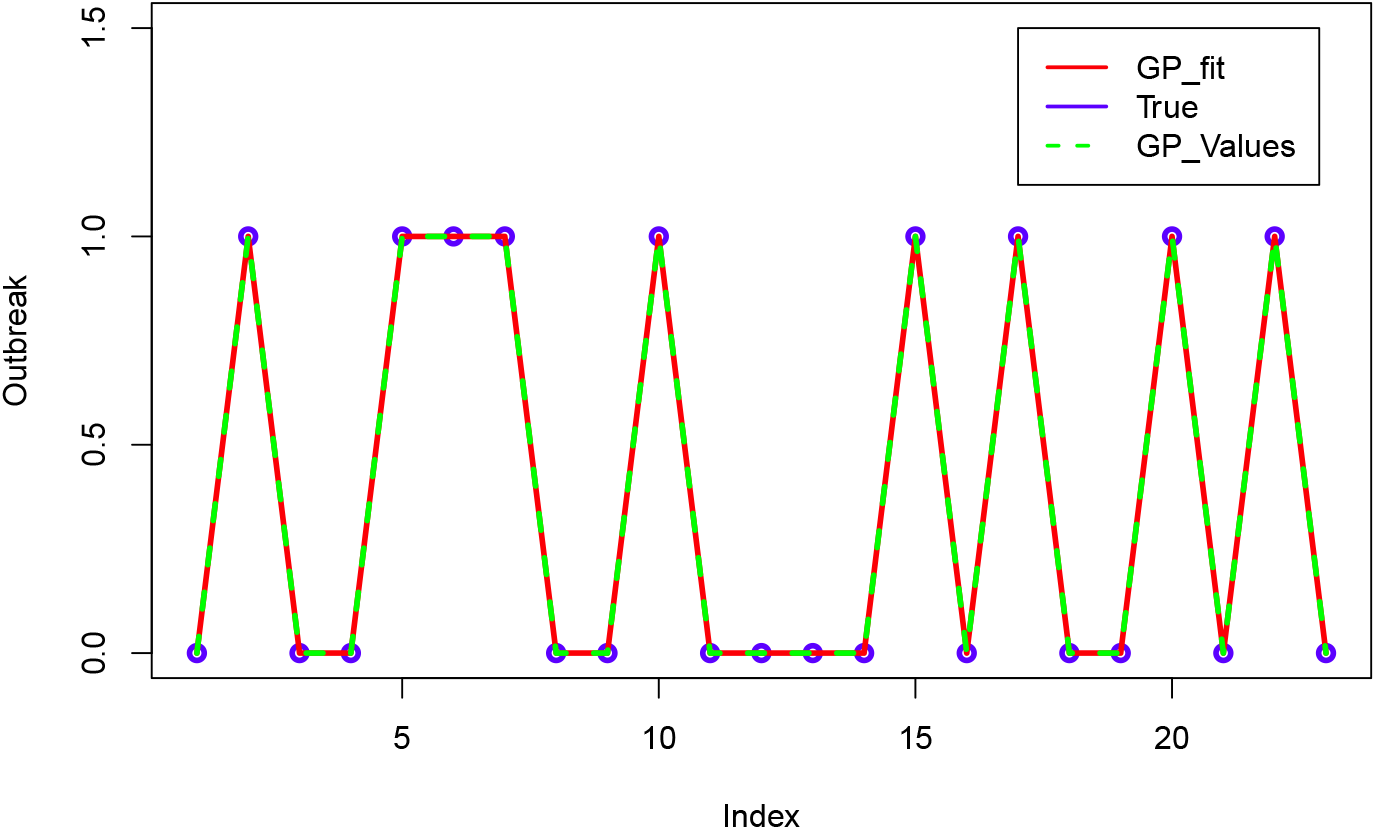
Fitting results from Gaussian Processes

**Figure 6:**
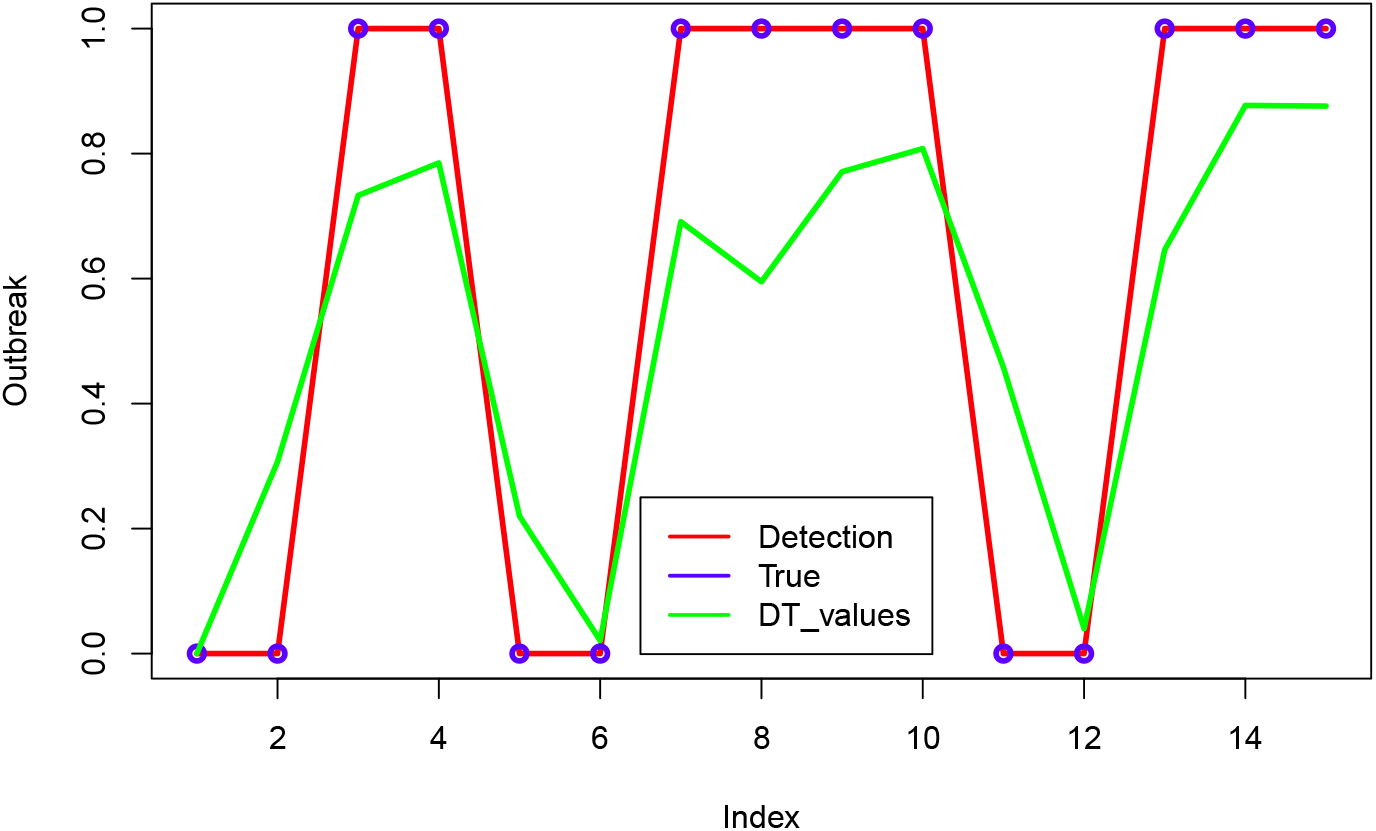
Detection results from random decision trees

**Figure 7:**
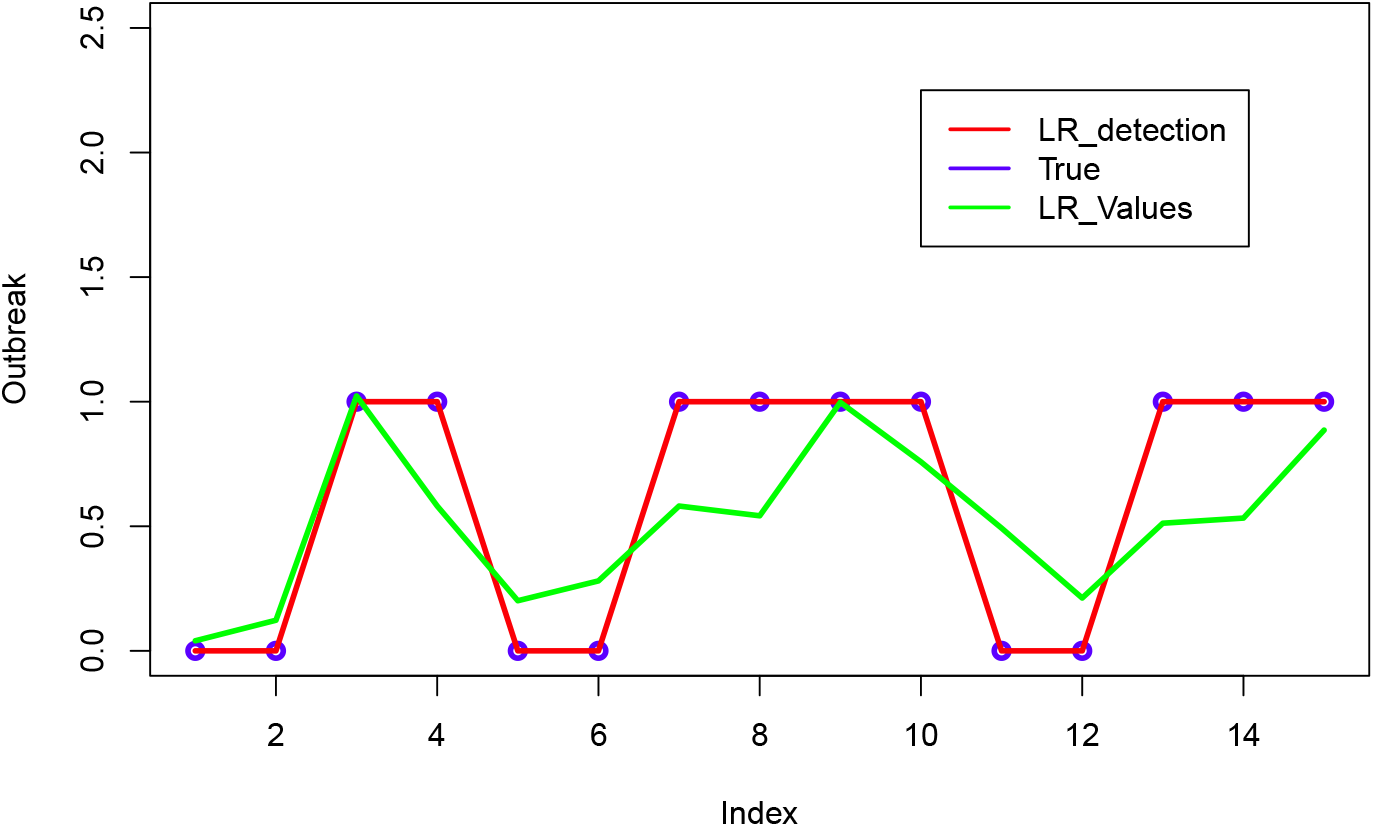
Detection results from logistic regression

**Figure 8:**
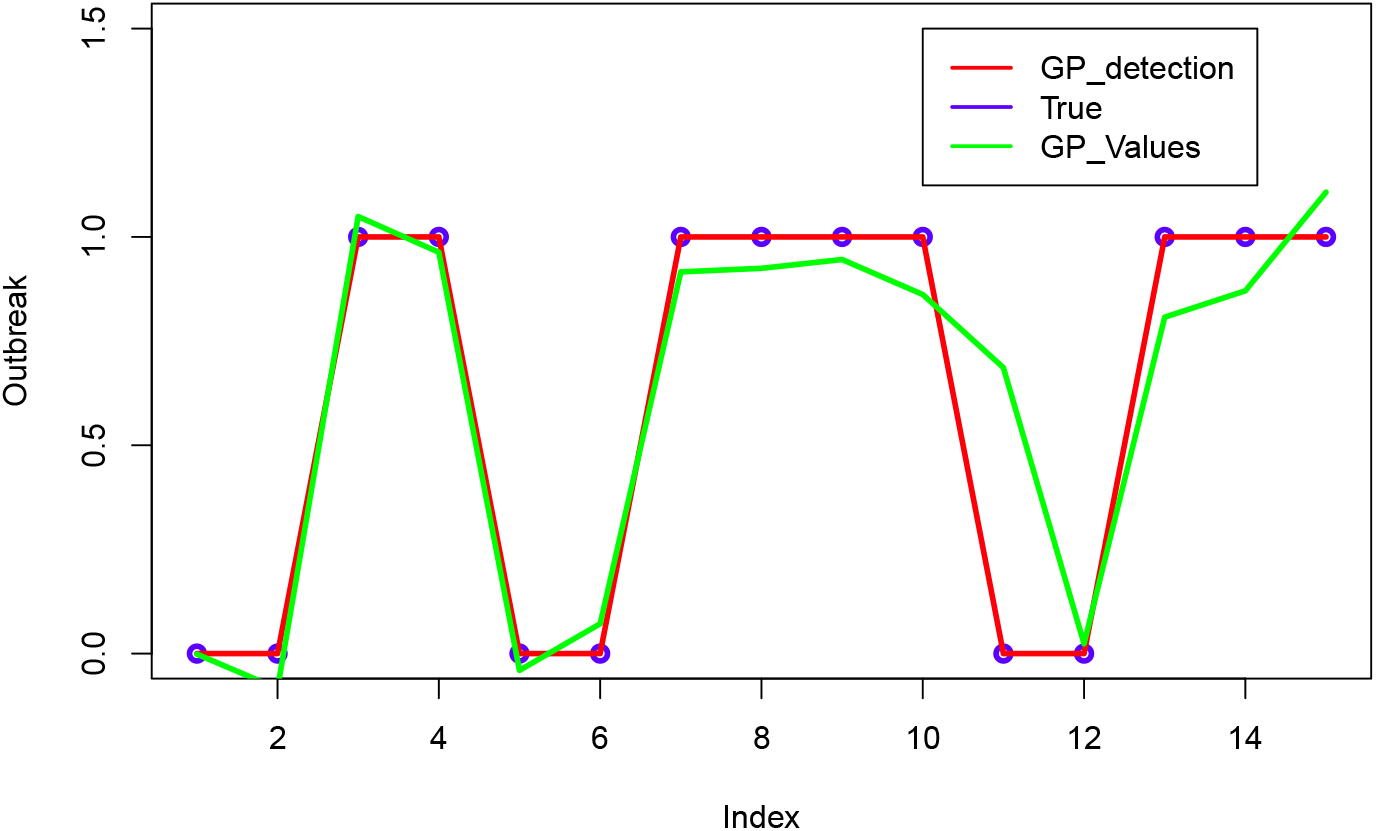
Detection results from Gaussian Processes

Table 1 shows the comparison of the test results from compared models. Errors are false positive when there is no outbreak but the methods predicts one and false negative when there is an outbreak but model does not detect one. Certainly, false negative can be considered more serious in detection problems. The methods proposed in this study give 100% accuracy of detection.

**Table 1:** Testing data sample and detection results

| Maxtemp | Mintemp | hum | rf | pos | pF | Outbreak | ANN* | SVM* | DT | LR | GP |
| --- | --- | --- | --- | --- | --- | --- | --- | --- | --- | --- | --- |
| 35 | 25 | 58.14 | 0 | 5221 | 110 | 0 | 0 | 0 | 0 | 0 | 0 |
| 35 | 27 | 65.13 | 10.09 | 7452 | 498 | 0 | 0 | 0 | 0 | 0 | 0 |
| 33 | 24 | 67.42 | 4.29 | 11856 | 504 | 1 | **0** | 1 | 1 | 1 | 1 |
| 32 | 27 | 83.19 | 15.63 | 10598 | 614 | 1 | 1 | 1 | 1 | 1 | 1 |
| 34 | 26 | 81.73 | 11.34 | 8432 | 593 | 0 | **1** | 0 | 0 | 0 | 0 |
| 36 | 26 | 64.39 | 4.12 | 9230 | 498 | 0 | 0 | 0 | 0 | 0 | 0 |
| 35 | 24 | 53.87 | 0 | 10745 | 453 | 1 | **0** | **0** | 1 | 1 | 1 |
| 35 | 27 | 84.97 | 14.55 | 10639 | 313 | 1 | 1 | 1 | 1 | 1 | 1 |
| 34 | 24 | 85.48 | 7.91 | 11823 | 549 | 1 | 1 | 1 | 1 | 1 | 1 |
| 34 | 25 | 81.2 | 12.08 | 11276 | 443 | 1 | 1 | 1 | 1 | 1 | 1 |
| 36 | 24 | 64.58 | 0.3 | 10389 | 591 | 0 | 0 | 0 | 0 | 0 | 0 |
| 32 | 22 | 64.3 | 0.64 | 8543 | 365 | 0 | 0 | 0 | 0 | 0 | 0 |
| 36 | 24 | 69.23 | 0 | 10545 | 341 | 1 | 1 | 1 | 1 | 1 | 1 |
| 38 | 22 | 26.94 | 0 | 10631 | 560 | 1 | 1 | 1 | 1 | 1 | 1 |
| 38 | 29 | 77.43 | 16.67 | 11732 | 462 | 1 | **0** | 1 | 1 | 1 | 1 |

\*. Results from [1], in bolds are errors

## 5 Conclusions and Discussion

Based on the sample data used, random decision trees, logistic regression, and Gaussian Processes provide promising results. All of the models were able to detect malaria outbreak with 100% accuracy on the test data. There are a few key points to be considered for each model:

- Decision trees should be randomized. Reasonable data amount should be used to avoid excessive recursive evaluations. In fact, with shorter training and test datasets, random decision trees can be used within rolling horizon framework. The advantage of the method is that no special treatment of dimensionality reduction on explanatory variables is needed.
- Logistic regression needs careful determination of the distribution family and subset of the variables. In this problem Poisson family with log link function worked best with a subset of 3 variables. Advantage of this method is its simplicity.
- Gaussian Process regression also needs determination of a subset of highly impacting factors. This is understandable within detection or classification problem. Also variables may be needed to be transformed for GPs.
- Overall, each method is more appropriate for regression problem. We had to determine threshold values to detect outbreak. Since these are determined in training phase, it does not negatively impact the decision making in testing phase.

For future work, models can be compared using similar large datasets for stronger conclusions.

## Supporting information

Dataused

## Acknowledgments

This study is supported by the Center for Connected Multimodal Mobility (*C*^2^*M*^2^) (USDOT Tier 1 University Transportation Center) Grant headquartered at Clemson University, Clemson, South Carolina, USA. The authors would also like to acknowledge U.S. Department of Homeland Security (DHS) Summer Research Team Program Follow-On, FY19 US Department of Education MSEIP Grant Award P120A190061, and National Science Foundation (NSF, No. 1719501, 1436222, and 1400991) grants. Any opinions, findings, conclusions or recommendations expressed in this study are those of the author(s) and do not necessarily reflect the views of (*C*^2^*M*^2^), USDOT, DHS, U.S. Department of Education, or NSF and the U.S. Government assumes no liability for the contents or use thereof.

