## Supplementary material for "Malaria Outbreak Detection with Machine Learning Methods": Dataused

| id | maxtemp | mintemp | humidity | rainfall | positive | pf | outbreak |
|---|---|---|---|---|---|---|---|
| 1 | 29 | 18 | 49.74 | 0 | 2156 | 112 | 0 |
| 2 | 34 | 23 | 83.27 | 15.22 | 10717 | 677 | 1 |
| 3 | 40 | 23 | 50.74 | 0 | 1257 | 127 | 0 |
| 4 | 34 | 24 | 59.16 | 9.06 | 4198 | 211 | 0 |
| 5 | 34 | 27 | 73.23 | 0 | 11808 | 712 | 1 |
| 6 | 31 | 24 | 88.77 | 41.4 | 10881 | 648 | 1 |
| 7 | 33 | 24 | 77.94 | 23.88 | 8830 | 459 | 1 |
| 8 | 31 | 24 | 84.57 | 11.15 | 9693 | 482 | 0 |
| 9 | 36 | 24 | 53.4 | 2.12 | 9310 | 549 | 0 |
| 10 | 32 | 23 | 57.5 | 0 | 13154 | 838 | 1 |
| 11 | 34 | 18 | 59.4 | 0 | 2197 | 136 | 0 |
| 12 | 42 | 24 | 49.43 | 2.19 | 3362 | 213 | 0 |
| 13 | 45 | 32 | 34.74 | 0.38 | 416 | 26 | 0 |
| 14 | 43 | 28 | 69.07 | 4.65 | 7514 | 410 | 0 |
| 15 | 33 | 23 | 80.97 | 6.92 | 10990 | 390 | 1 |
| 16 | 32 | 24 | 87.32 | 11.92 | 6536 | 338 | 0 |
| 17 | 40 | 27 | 63.97 | 0 | 11169 | 776 | 1 |
| 18 | 39 | 25 | 47.52 | 0 | 8131 | 312 | 0 |
| 19 | 36 | 26 | 72.78 | 3.54 | 5138 | 213 | 0 |
| 20 | 31 | 23 | 73.35 | 4.97 | 10659 | 612 | 1 |
| 21 | 30 | 23 | 86.81 | 7.21 | 9041 | 418 | 0 |
| 22 | 30 | 22 | 78.8 | 3.12 | 11265 | 404 | 1 |
| 23 | 33 | 22 | 73.71 | 1.75 | 9233 | 212 | 0 |
| 24 | 35 | 25 | 58.14 | 0 | 5221 | 110 | 0 |
| 25 | 35 | 27 | 65.13 | 10.09 | 7452 | 498 | 0 |
| 26 | 33 | 24 | 67.42 | 4.29 | 11856 | 504 | 1 |
| 27 | 32 | 27 | 83.19 | 15.63 | 10598 | 614 | 1 |
| 28 | 34 | 26 | 81.73 | 11.34 | 8432 | 593 | 0 |
| 29 | 36 | 26 | 64.39 | 4.12 | 9230 | 498 | 0 |
| 30 | 35 | 24 | 53.87 | 0 | 10745 | 453 | 1 |
| 31 | 35 | 27 | 84.97 | 14.55 | 10639 | 313 | 1 |
| 32 | 34 | 24 | 85.48 | 7.91 | 11823 | 549 | 1 |
| 33 | 34 | 25 | 81.2 | 12.08 | 11276 | 443 | 1 |
| 34 | 36 | 24 | 64.58 | 0.3 | 10389 | 591 | 0 |
| 35 | 32 | 22 | 64.3 | 0.64 | 8543 | 365 | 0 |
| 36 | 36 | 24 | 69.23 | 0 | 10545 | 341 | 1 |
| 37 | 38 | 22 | 26.94 | 0 | 10631 | 560 | 1 |
| 38 | 38 | 29 | 77.43 | 16.67 | 11732 | 462 | 1 |
